# Evaluation of the efficacy of simethicone in the abdominal ultrasound and pancreas evaluation on dogs

**DOI:** 10.1101/2021.12.01.470744

**Authors:** L. F. C. Garcia, B. H. Cottar, M. Rodacki, M. M. Gonçalves, V. Grendel, L. Albrecht, S. H. Weber, U. I. Tasqueti

## Abstract

Intestinal gas results in low quality imaging in abdominal ultrasound in domestic animals. On dogs, suggested preparation protocols are varied and low studied. The aim of this work was comparing the efficacy between two preparations in improving the complete abdominal ultrasound exam and pancreas preview. 40 dogs were enrolled in this study. They were separated in two groups (NS diet: fasting; WS diet: fasting added to simethicone). The pancreas images were taken from left lateral decubitus. A score of one to three (1: bad, 2: fair and 3: good) measured separately the quality of the complete exam, left and right limbs and pancreatic body. In both treatments, there were results above 50% of good images of the complete exam and right limb of the pancreas. Otherwise, there were results above 90% of bad images of the right limb and pancreatic body. According to the fact of there is no statistically significant difference between the diets, as well submitted animals to the NS diet as the WS diet, all the results obtained from this work (bad or good images) could be acquired from any of both diets. The good abdominal exam preview as well submitted animals to the simethicone as those without the medication refute this medicine requirement to the abdominal ultrasound exam, but not to the evaluation of the left limb and pancreatic body.

## RESUMO

Gases intestinais resultam em imagens de má qualidade na ultrassonografia abdominal de animais domésticos. Em cães, os protocolos de preparo sugeridos são variados e pouco estudados. O objetivo desse trabalho foi comparar a eficácia entre dois preparos na melhora da avaliação ultrassonográfica abdominal completa e visualização do pâncreas. Quarenta cães foram envolvidos nesse estudo. Eles foram separados em dois grupos (dieta NS: Jejum; dieta WS: Jejum associado à simeticona). As imagens do pâncreas foram obtidas a partir do decúbito lateral. Um escore de um a três (1: ruim; 2: razoável; 3: bom) foi usado para avaliar separadamente a qualidade do exame completo, lobos esquerdo e direito e corpo pancreático. Em ambos os tratamentos, houve mais que 50% de boas imagens obtidas a partir do exame abdominal completo e lobo pancreático direito. Por outro lado, houve mais que 90% de imagens ruins do lobo direito e corpo pancreático. De acordo com o fato de não haver diferença estatisticamente significativa entre as dietas, tanto animais submetidos à dieta SS como à dieta CS, todos os resultados obtidos a partir desse trabalho foram obtidas (sejam imagens boas ou ruins) puderam ser adquiridas a partir de ambas as dietas. A boa avaliação abdominal completa tanto em animais submetidos como aqueles sem a medicação refuta o requerimento medicamentoso para a avaliação ultrassonográfica, mas não para a avaliação dos lobos esquerdo e corpo pancreático.

**Palavras Chave:** Canino; imagem; gás intestinal; artefato.

## INTRODUCTION

Ultrasound is a very well accepted and successful veterinary method for anomalies identification. It is understood that the first record of a clinical sonography investigation goes back to sixth decade, when pregnancy on sheep was diagnosed for the first time (LINDAHL, 1966). Since this period, ultrasonography had an increasing relationship with veterinarians. In view of a large amount of benefits credited to this method, it should be used as a vital tool to the veterinarian. Notwithstanding, among such benefits, there is a succession of reported complications along an ultrasound exam. After knowing such situation, it is highly necessary to understand and comprehend all the different images this exam could provide. Among the frequent situations faced by this professional, there are the artefacts produced by the gases (OHLERTH and O’BRIEN, 2007).

Some studies presume the importance added to the prior preparation of this professional to distinguish a true image from a false report (LARSON, 2016). In general, with exception of emergencies, imaging diagnosis services require an easier preparation for abdominal ultrasound examination, which involves water ingestion for the urinary bladder filling and better pancreas imaging. A wide abdominal trichotomy caudal to the xiphoid process, ventral to the epaxial muscles and cranial to the pelvis is also recommended (CARVALHO, et al., 2008). Among the application of different methods to assist the abdominal imaging for humans or animals, one of them still under discussion by different authors: the application of antiphysetic medicine (CHANG, et al., 2014).

The simethicone (dimethylhexane) is featured by an association of dimethicone and silicone dioxide. Due to the silicone low tension surface effect, it is capable of promote intestinal gas bubbles agglutination, resulting in an easier dispersion. According to this reaction, this medicine is recommended to the intestinal gas volume reduction and support an ultrasound examination (GE, et al., 2006). It has been reported, by human endoscopy studies in 2006, that there is a clear differentiation between evaluated images from small bowel accomplished by the assistance of simethicone from the others without any medicine support (GE, et al., 2006).

Many ultrasound and endoscopy studies largely resort to this drug as an essential element, although there is no consensus about the ideal drug preparation for animals. Even for humans, the ideal dosage, volume and frequency for this premedication still unsure (CHANG, et al., 2014). Authors evidence the relationship of such variability with equipment technology development which, in given situation, requires different period drug treatment while in other situations, the application becomes dispensable (CHANG, et al., 2014; GRANGER, et al., 2015). It was observed that fasting does not represent a significant fostering to the abdominal evaluation of different organs on dogs (GARCIA; FROES, 2014). Otherwise, the efficacy of anti-foaming agents, like simethicone, still uncertainty.

## MATHERIALS AND METHODS

### Study Group

This work was accomplished under the Research Ethical Committee registration number 0927 – first version. Altogether, 40 dogs were selected to this study. 24 (60%) of these were female and 16 (40%) were male. Due to the animals were presented to the Hospital Unit for Company Animals because of different diseases, it was necessary the development of exclusion criterions, which prevented erroneous results to this work. The exclusion criterion were: diarrhea, vomit, clinical or surgical emergency, statement of any gastrointestinal condition and presence of any kind of peritoneum effusion (MORTIER, et al., 2016). There are no studies which report imaging complication due to age or weight. However, under 3 months animals did not participate of this study due to all of them have arrived the hospital with diarrhea.

### Technique

It was made a division in two groups compound by 20 individual each. The first group received a treatment compound by solid fasting 8 hours before the ultrasound examination (NS diet). The second group was submitted to a treatment concerning a solid fasting 8 hour before the ultrasound examination added to the administration of simethicone, 4 mg/kg, three times a day along 48 hours before the ultrasound examination (WS diet). Owners of all animals were properly instructed about moment and ways of administration.

A group compound by two image diagnosis resident veterinaries was previously selected to the achievement of the research image evaluation. All of them have received exactly the same evaluation terms. The appraisal was developed by only one ultrasound device model Esaote MyLab40® and a micro convex probe on frequency of 5 MHz. The development of the imaging was elapsed in a low luminosity, distraction and stress environment.

### Image Analysis

Two evaluations compound by a score of one to three (1 – Bad, 2 – Fair and 3 – Good) was applied to the imaging evaluation (GE, et al., 2006). The first one was referred to the quality of the routine complete ultrasound examination, which was compound sequentially by urinary bladder, prostate (male), uterus (female), left kidney, left adrenal gland, spleen, liver, gall bladder, stomach, pancreas, right kidney and right adrenal gland; the second evaluation measured the pancreatic body, left and right limbs imaging quality. The score was solely based on the relationship between the gas volume and the imaging commitment by following remarks: 1 – Bad: large gas amount preventing a part or all the examination; 2 – Fair: moderate gas amount significantly allowing the examination; 3 – Good: a few or no gas presence preventing the examination. Artefacts presence were no evaluated in this study.

The routine ultrasound examination was performed with experimental units on dorsal decubitus. Pancreas evaluation was performed by a change to left lateral decubitus. All the acquired images were evaluated by one of the two ultrasonographers by a blind random assortment. The professional had no information about which diet each experimental unit was submitted.

### Data Analysis

The Kolmogorov-Smirnov test, featured by the agreement degree mensuration between sample values of different group distribution, was considered the most indicated test to this study development. (RAZALI and WAH, 2011; SHIROUCHAKI, et al., 2014).

## RESULTS

The obtained values from complete abdominal imaging represented by the figure 1 did not present a statistically significant difference (Dmax 0,25 < KS tab 0,34). From all the dogs submitted to the NS diet, two experimental units (10%) propitiated a bad imaging quality (figure 1A), 11 (55%) fair (figure 1B) and seven (35%) good (figure 1C). From all the dogs submitted to the WS diet, one (5%) propitiated a bad imaging (figure 1A), eight (40%) fair (figure 1B) and 11 (55%) good (figure 1C).

**Figure 1.**
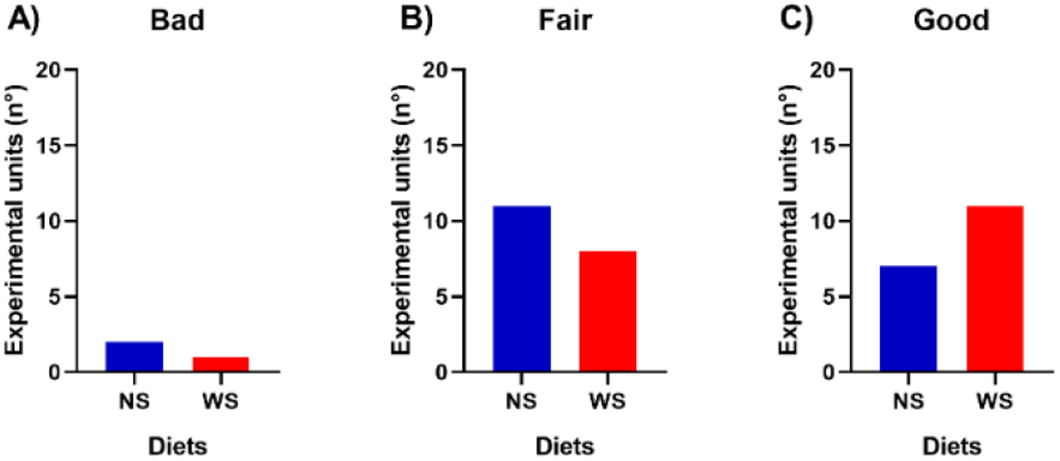
Pretreatment with simethicone o not foster the complete abdominal ultrasound evaluation. After the agreement of the owners, the dogs were submitted to the simethicone-based diet, 4 mg/kg, three times a day during three days (WS diet) or placebo diet (NS diet) before the imaging evaluation. At the day of the evaluation, the animals were conducted to the Imaging Department after abdominal trichotomy. They were positioned in dorsal decubitus and the routine abdominal imaging was done by an experienced veterinary. At the ending of the evaluation, the veterinary scored the quality of the obtained images as bad **(A)**, fair **(B)** or good **(C)**. The professional did not know to which diet the animal was conducted. Dmax 0,25 < KS tab 0,34 (Kolmogorov-Smirnoff test).

The obtained values from the pancreatic right limb imaging represented by the figure 2 do not present a statistically significant difference (Dmax 0,0625 < KS tab 0,34). From all the dogs submitted to the NS diet, one experimental unit (5%) propitiated a bad imaging quality (figure 2A), nine (45%) fair (figure 2B) and 10 (50%) good (figure 2C). From all the dogs submitted to the WS diet, two (10%) propitiated a bad imaging (figure 2A), seven (35%) fair (figure 2A) and 11 (55%) good (figure 2C).

**Figure 2.**
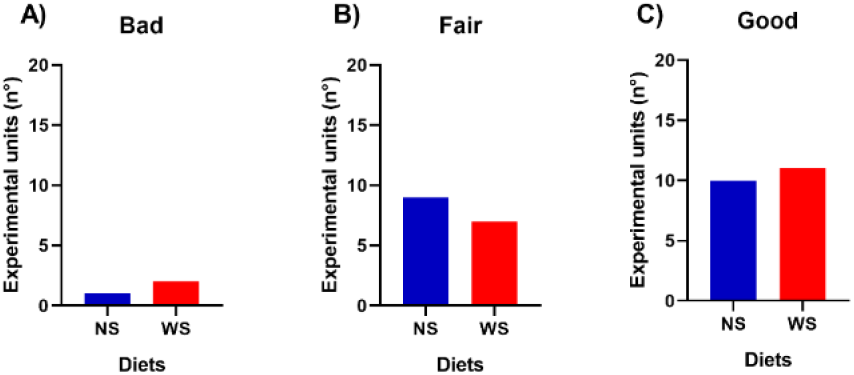
Pretreatment with simethicone do not foster the pancreatic right limb ultrasound evaluation. After the agreement of the owners, the dogs were submitted to the simethicone-based diet, 4 mg/kg, three times a day during three days (WS diet) or placebo diet (NS diet) before the imaging evaluation. At the day of the evaluation, the animals were conducted to the Imaging Department after abdominal trichotomy. They were positioned in left lateral decubitus and the imaging was done by an experienced veterinary. At the ending of the evaluation, the veterinary scored the quality of the obtained images as bad **(A)**, fair **(B)** or good **(C)**. The professional did not know to which diet the animal was conducted. Dmax 0,0625 < KS tab 0,34 (Kolmogorov-Smirnoff test).

The obtained values from the pancreatic body imaging represented by the figure 3 do not present a statistically significant difference (Dmax 0,125 < KS tab 0,34). From all the dogs submitted to the NS diet, 18 experimental units (90%) have propitiated a bad imaging quality (figure 3A) and only one (5%) a good imaging (figure 3C). From all the dogs submitted to the WS diet, all of them (100%) have propitiated a bad imaging quality (figure 3A). There was no fair result from the pancreatic body imaging (figure 2B).

**Figure 3.**
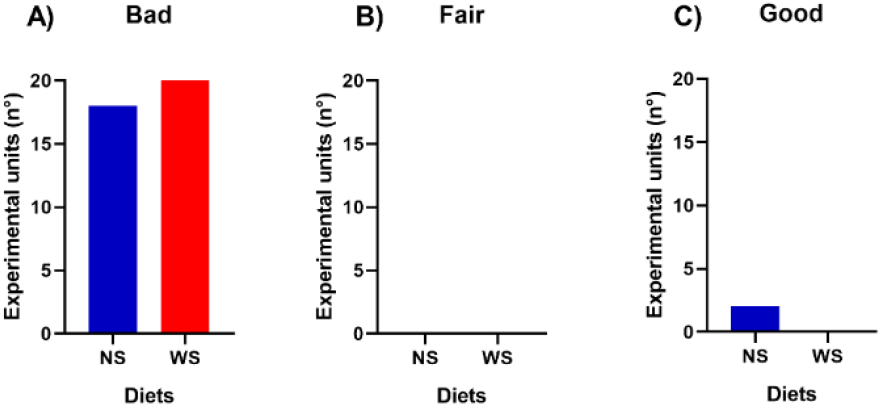
Pretreatment with simethicone do not foster the pancreatic body ultrasound evaluation. After the agreement of the owners, the dogs were submitted to the simethicone-based diet, 4 mg/kg, three times a day during three days (WS diet) or placebo diet (NS diet) before the imaging evaluation. At the day of the evaluation, the animals were conducted to the Imaging Department after abdominal trichotomy. They were positioned in left lateral decubitus and the abdominal imaging was done by an experienced veterinary. At the ending of the evaluation, the veterinary scored the quality of the obtained images as bad **(A)**, fair **(B)** or good **(C)**. The professional did not know to which diet the animal was conducted. Dmax 0,125 < KS tab 0,34 (Kolmogorov-Smirnoff test).

The obtained values from pancreatic left limb imaging represented by the figure 4 do not present a statistically significant difference (Dmax 0,0625 < KS tab 0,34). From all the dogs submitted to the NS diet, 19 experimental units (95%) propitiated a bad imaging quality (figure 4A) and only one (5%) a good imaging (figure 4C). From all the dogs submitted to the WS diet, all of them (100%) have propitiated a bad imaging quality (figure 4A). There was no fair result from the pancreatic left limb imaging (figure 4B).

**Figure 4.**
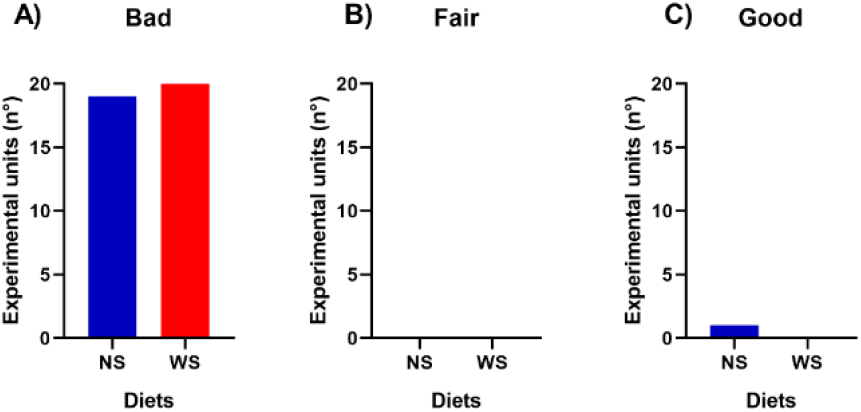
Pretreatment with simethicone do not foster the pancreatic left limb ultrasound evaluation. After the agreement of the owners, the dogs were submitted to the simethicone-based diet, 4 mg/kg, three times a day during three days (WS diet) or placebo diet (NS diet) before the imaging evaluation. At the day of the evaluation, the animals were conducted to the Imaging Department after abdominal trichotomy. They were positioned in left lateral decubitus and the abdominal imaging was done by an experienced veterinary. At the ending of the evaluation, the veterinary scored the quality of the obtained images as bad **(A)**, fair **(B)** or good **(C)**. The professional did not know to which diet the animal was conducted. Dmax 0,0625 < KS tab 0,34 (Kolmogorov-Smirnoff test).

## DISCUSSION

In 2014, a work involving 150 dogs demonstrated that fasting was a questionable pretreatment for the ultrasound evaluation of different organs (Garcia; Froes, 2014). Based on this work, we evaluated the efficacy of anti-foaming medicine on the ultrasound imaging on dogs.

The low rate of bad images obtained from the complete abdominal exam as well submitted animals to the NS diet composed by fasting (10%) as those submitted to the WS diet summed up to fasting added to simethicone (5%) suggest that fasting is enough for a routine abdominal imaging. The rate of fair images obtained (55% for NS diet; 40% WS diet) added to the rate of good images (35% for NS diet; 55% WS diet) from the same exam suggests that there is no difference between both treatments. Our results corroborate with different works. These researches demonstrated that the hindering of the evaluation of different organs could occur due to the presence of air or food on the intestinal tract (Barberet, et al., 2008). According to this fact, the evaluation of animals without a pretreatment could speed up the waiting time.

Pancreatic left limb offered the worst results in this work represented by 95% of NS diet and all the evaluations of the WS diet. It reveals a complete inefficacy of the current protocol to the ultrasound evaluation of this segment of the pancreas. The evaluation of the pancreatic body also demonstrated very expressive results which 90% of the group submitted to the NS diet and all the animals submitted to the WS diet provided bad quality images. In contrast to the results obtained from the left limb and pancreatic body, the evaluation of the right limb of this organ offered a low rate of bad quality images represented by 5% from NS diet and 2% from WS diet. These results disagree with some authors which affirm that the gas produced from stomach and duodenum might be the main causer agent of the bad quality imaging (Barberet, et al., 2008). Otherwise, it is in accordance with authors who affirm that the right limb of the pancreas is the easiest part of this organ to be viewed. (CARVALHO, 2004). The positioning of the pancreas, dorsomedial to the descending duodenum, evidences such possibility (CORTÉS, 2002). The good quality images obtained from the right limb of the pancreas suggest the generation of gas begins on the duodenum once this segment of the pancreas approaches the greater curvature and pylorus of the stomach (CORTÉS, 2002).

According to the fact of there is no statistically significant difference between the diets, as well submitted animal to the NS diet as the WS diet, all the results obtained from this work (bad or good images) could be acquired from any of both diets. There was also no report of difficulty or adverse effects related by the owners due to the administration of the treatments.

The pancreas is a very important organ to be evaluated. Acute or chronic pacreatitis, pancreatic edema, pseudocysts, abscess, nodular hyperplasia or even neoplasia are diseases that could be observed by a well implemented ultrasound imaging (LARSON, 2016). Such problems could reveal a base disease or provide us information about the evolution of different disorders. According to it, it is extremely important to evaluate the dog pancreas with speed and precision.

The bad rate of the pancreatic body and left limb imaging obtained from animals submitted or not to the medication demands for alternative methods such like decubitus or incidence change. Different methods applied in humans, such like Simethicone Water Rotation, represent satisfactory results in all pancreas imaging (ISHIGAMI, 2014). This condition inspires new studies about optional techniques on dogs. According to Nakamura, et al. (2015), pancreatic insulinoma is a harmful cancer which needs a different technique to be identified by the ultrasound exam. It is characterized by the intravenous application of perfublatane microbubble contrast agent. Such technique should not be applied in all routine ultrasound exam duo to it invasive characteristic. But, facing any pancreatic threat, it might be considered.

Besides the aim of this work was not to evaluate the relationship between dog size and the quality of the ultrasound preview, two cases involving giant dogs showed interesting results, they were the unique animals which resulted in a good pancreatic body imaging. According to Larson (2016), animals with a deep chest conformation propitiate a right lateral intercostal window, which is extremely helpful.

## CONCLUSIONS

The good rate of complete abdominal exam preview as well submitted animals to the simethicone as those without the medication suggests that this medicine is not necessary for abdominal imaging. Such situation may reduce the waiting time for ultrasound exams in animal hospitals.

## ACKNOWLEDGMENT

Thanks to the Hospital Unit for Companion Animals for all the offered infrastructure and support. The good development of this project has only occurred by the valorous help from the Hospital team.

