## Supplementary figures and images for "Evaluation of the efficacy of simethicone in the abdominal ultrasound and pancreas evaluation on dogs"

### Supplemental material 1

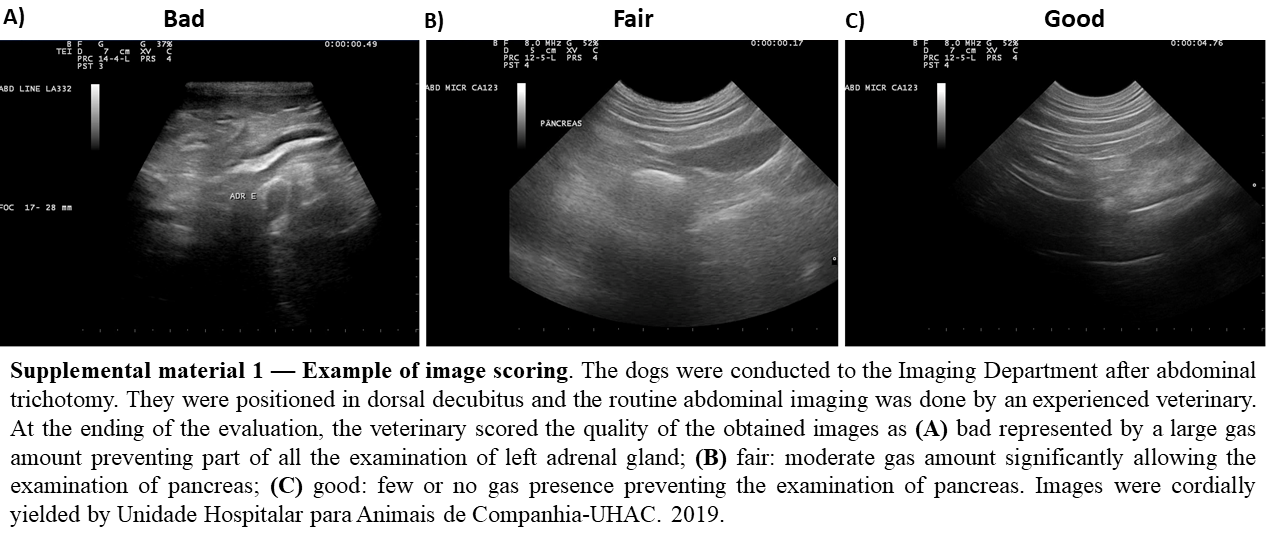
